# Long-term tree population growth can predict woody encroachment patterns

**DOI:** 10.1101/2024.04.07.588197

**Authors:** Robert K. Shriver, Elise Pletcher, Franco Biondi, Alexandra K. Urza, Peter J. Weisberg

## Abstract

Recent increases of woody plant density in dryland ecosystems around the world are often attributed to land use changes such as increased livestock grazing and fire suppression, or to climatic trends driven by increasing atmospheric carbon dioxide^1,2^. While such changes have undoubtedly impacted ecosystem structure and function, the evidence linking them to woody encroachment is mixed and demographic processes underlying changes in woody plant abundance require further consideration^3^. After examining tree age structures from woodlands across the interior western USA using demographic models, we find little evidence of widespread increases in per-capita tree establishment rates following 19^th^ century Euro-American settlement. Woodlands dominated by young trees have often been cited as evidence of woody encroachment driven by a number of anthropogenic processes, but we demonstrate they can also be accurately predicted by a null model including only steady long-term tree population growth. Contrary to common interpretations, we show that tree establishment rates in the last century have mostly declined, rather than increased, and in fact they are currently at their lowest rates since at least 1600 CE.

## Main Text

One of the most evident ecological changes in many dryland ecosystems over the last 200 years has been increasing density and distribution of woody vegetation^1,2^. Collectively referred to as woody encroachment or expansion, such changes have significantly impacted ecosystem processes and services, including global carbon cycling and hydrology^4,5^. Evidence of increasing woody plant density and cover has been found in several archives, including packrat middens, tree-ring establishment records, historical photos, modern field observations, and more recently, remote sensing^6,7^. In the western USA, tree-ring-derived age structures show dry woodlands, particularly pinyon-juniper woodlands, dominated by trees that largely established after the mid-1800’s CE^8^. Because this timing also coincides with the period of widespread Euro-American settlement in the region, the increase in establishment has usually been attributed to anthropogenic mechanisms, such as wildfire suppression and livestock grazing, or to changes in environmental conditions that coincide with recent Euro-American settlement (e.g. increasing atmospheric CO_2_)^6^. However, evidence supporting these hypotheses has generally not been conclusive or pervasive enough to explain regionwide increases in tree establishment^1,3^. For example, wildfire suppression and exclusion are often cited as a mechanism underlying increases in woodland density, but data on wildfire regimes indicate many pre-settlement pinyon-juniper woodlands were likely characterized by infrequent, high-severity fires, rather than frequent, low-severity fires that would have regularly reduced young tree density^3^. Similarly, impacts of overgrazing by domestic livestock are also commonly invoked to explain increasing woodland density, but sites with little livestock grazing history have seen similar increases in tree establishment^9,10^. As a result, directly attributing increasing woody plant density to specific external drivers remains a challenge^1,3,11^.

Demographic rates, and in particular the processes that control plant establishment (seed production, germination, and survival at early life stages), are critical in controlling rates of plant population growth ^12^. If hypothesized changes in abiotic or biotic conditions were responsible for a sudden increase in woody plant density, we would expect this to be associated with an increase in the per-capita establishment rate (new trees established per existing tree) sufficient to account for the observed increases in woody plant density^13^. For example, if high-frequency, low-severity fires previously killed seedlings and maintained low tree density, as has been observed in some ponderosa pine-dominated ecosystems of the southwest USA^14^, then fire exclusion would lead to a sudden increase in seedling survival and thus per-capita establishment rates. However, population growth is a non-linear, multiplicative process and thus rapid increases in population density and establishment over time are possible even if the underlying demographic rates remain constant, unchanged by environmental conditions (Fig. 1). Multiplicative population growth could create the false appearance of a sudden, recent increase in woody plant density driven by changing environmental conditions, especially when a population experiences prolonged periods of growth after colonizing new habitat or recovering from past disturbance^15^. This component of vegetation change is often overlooked but can be readily quantified using population models. Thus, determining the demographic and population processes behind increasing woody plant density is essential to understanding its causes^16^.

**Figure 1.**
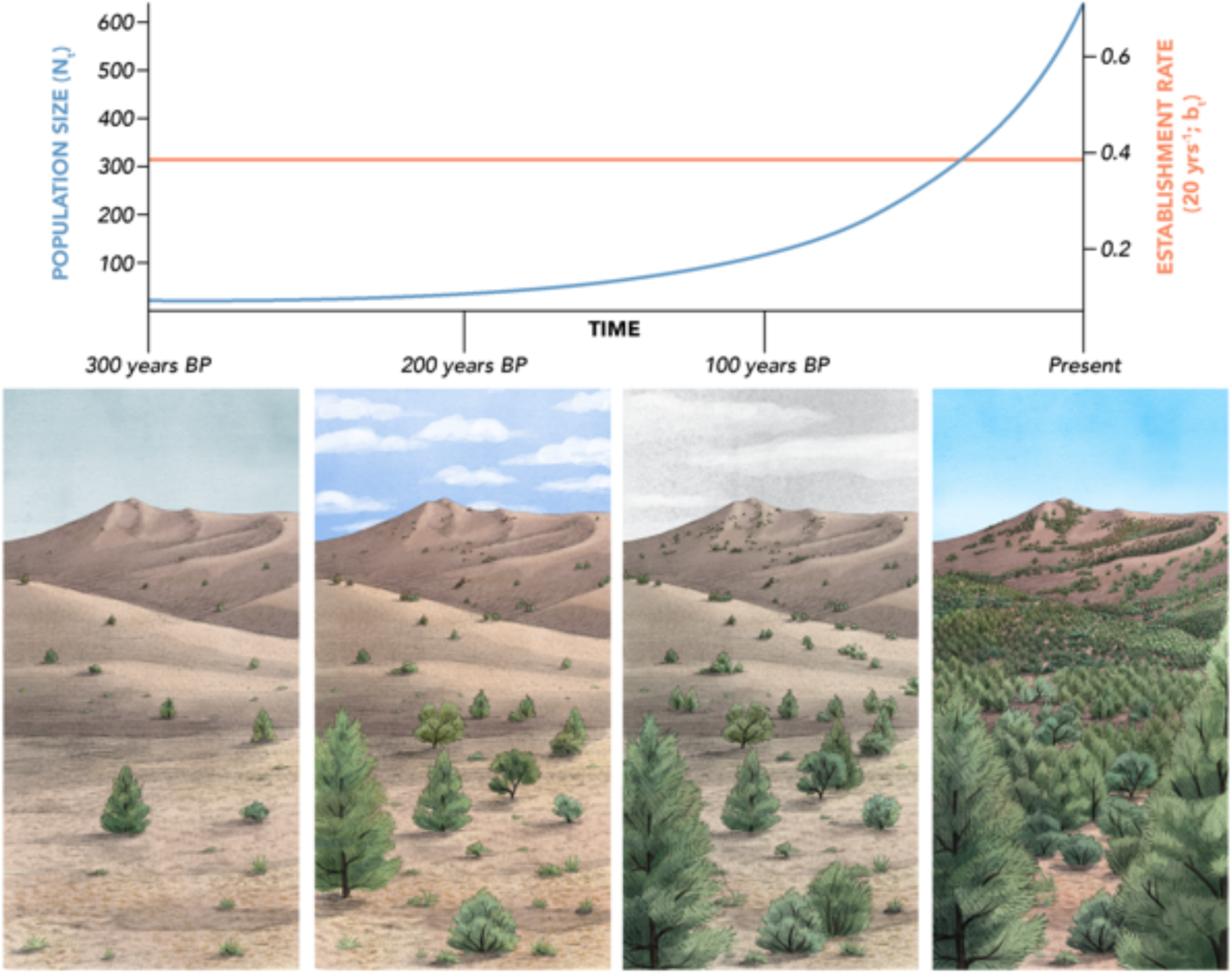
Increasing establishment under a constant establishment rate. Because new establishment (*B*_*t*_) is the product of both population size (*N*_*t*-1_) and per-capita establishment rate (*b*_*t*_; *B*_*t*_ = *b*_*t*_*N*_*t*-1_) in a growing population the amount of new establishment will increase over time even if the underlying establishment rate is constant. One could view the rapid increase in establishment and population size as evidence of a change point and link the increase in establishment to an external driver. However, analysis of the underlying demographic mechanisms will find that no change in per-capita establishment rates (orange) has occurred. Increased establishment is instead driven by the reproductive potential of larger populations. BP indicates “before present”. (Illustration by Alex Boersma)

Pinyon-juniper woodlands are among the most widespread forest types in the US, covering over 40 million hectares, and the most abundant remaining old-growth forest type in the US, representing nearly one-third of all old growth forest area^17^. While increases in density of pinyon-juniper woodlands have been widely noted, the causes and management of increasing woodland density are often controversial^3,18^. Despite a lack of conclusive evidence, the notion that increasing tree density and extent in western woodlands originated from Euro-American settlement remains prevalent in the scientific and land management literature. Such interpretation has motivated extensive vegetation treatments, usually framed as restoration efforts, to remove pinyon-juniper woodlands^19–21^. At the same time, these woodlands have also experienced widespread mortality that has been attributed to climate change, either directly as drought stress ^22^ or indirectly as a result of insect outbreaks and increased wildfire activity^23^.

One key aspect missing from previous interpretations of increasing woodland density is a quantification of the role of demographic and intrinsic population processes^16^. In this study, we used published age structure data from 29 populations at 23 sites in the Great Basin and Colorado Plateau, western USA (Fig. S1; Table S1), to calculate empirical per capita-establishment rates (hereafter “establishment rates”) prior to, during, and after Euro-American settlement. We then evaluated the impact that changing establishment rates had on observed woodland density increases by comparing these increases to a null model that uses only constant establishment rates. Extensive age structure data documenting woody encroachment from stands that predate settlement are rare, making pinyon-juniper woodlands an ideal case study.

### Tree establishment rates since 1600

Age structure data indicated a regional trend of increasing pinyon-juniper tree establishment and density during the late 1800s and early 1900s CE, with more than half of existing trees establishing after 1850 CE in all but two populations (Fig. 2A). We found, however, that establishment rates between 1860-1900 CE, the period of recent Euro-American settlement, generally fell within the historic range of variability (Fig. 2B, S2), suggesting that recent increases in establishment and density may be explained by long-term tree population growth rather than anthropogenic or environmentally-driven changes to underlying establishment rates.

**Figure 2.**
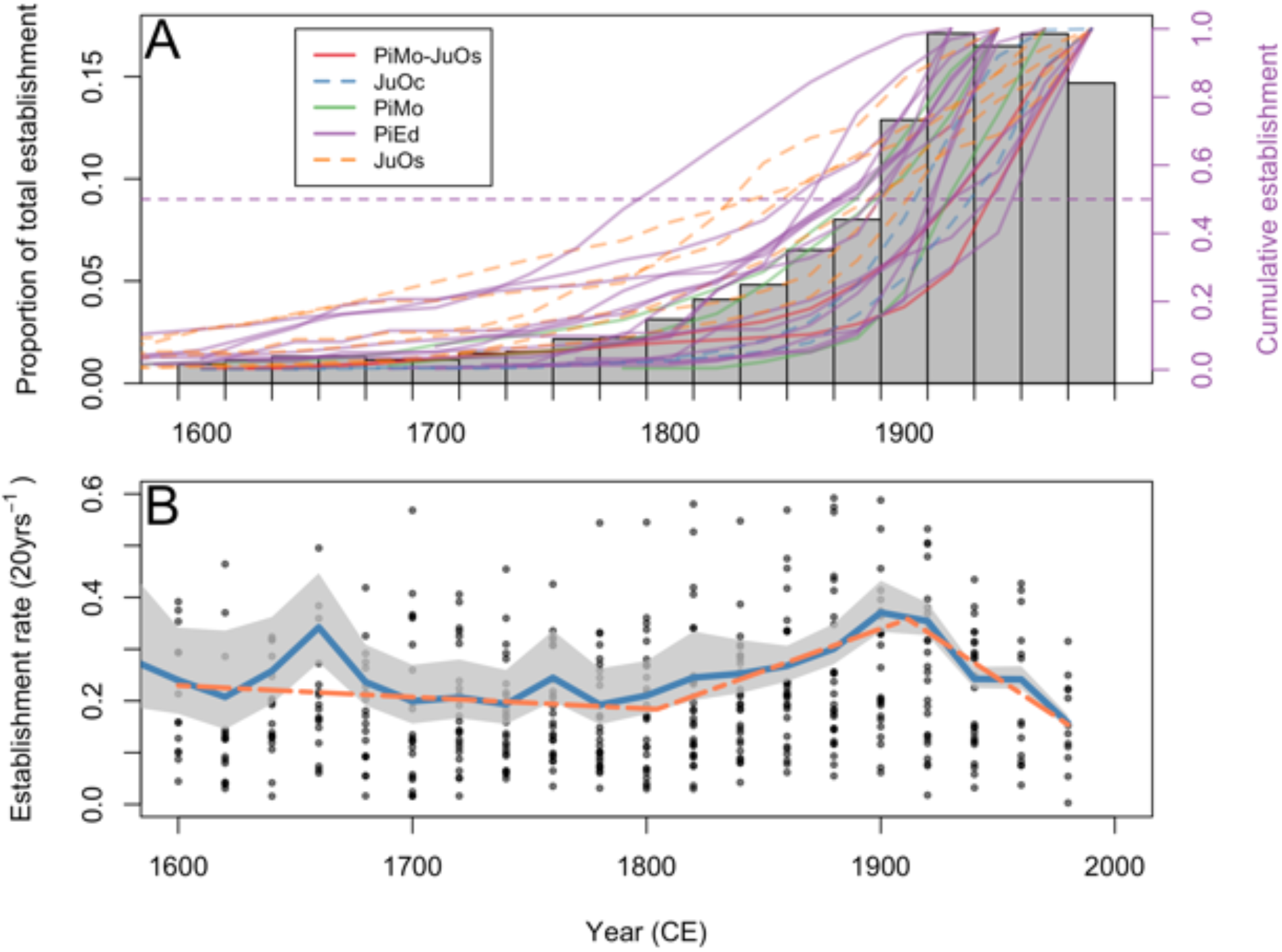
Tree establishment rates since 1600 CE. (A) Proportion of total observed establishment occurring in each 20-year interval averaged across 29 populations and cumulative observed establishment in each population broken down by species. Dashed horizontal line indicates 50% of total. Species acronyms in legend are mixed stands dominated by *Pinus monophylla*, with a minor *Juniperus osteosperma* component (PiMo-JuOs), *Juniperus occidentalis* (JuOc), *Pinus monophylla* (PiMo), *Pinus edulis* (PiEd), and *Juniperus osteosperma* (JuOs). (B) Per-capita establishment rates for each population (points), 20-year average (blue) and 95% credible interval (grey), and the best fit segmented regression (dashed orange). The entire timeseries dataset is shown in Fig. S2.

Prior to 1600 CE, average establishment rates were highly uncertain and variable, fluctuating between 0.14 and 0.48 (0.005-1.64 95% CI) average per-capita establishment rate per 20-year interval (Fig. S2). Following approx. 1600 CE average establishment rates became more certain. Shortly after 1600 CE establishment rates dropped to 0.21 (0.17-0.35 95% CI) per 20-year interval, but then increased to 0.34 (0.28-0.45 95% CI) in 1660 CE, before briefly stabilizing at ∼0.20 between 1680 and 1740 CE (Fig. 2B). In the late-1700s CE, average establishment rates gradually increased until 1920 CE. The results of segmented regression further corroborated this modest upward trend in establishment (Table S2). Although the total number of establishing trees per 20-year period tripled from the 1820 to 1900 CE, we found only a modest increase in average per-capita tree establishment rates from 0.24 in 1820 CE (0.2-0.33 95% CI) and to 0.37 in 1900 CE (0.33-0.44 95% CI) (Fig. 2A,B). Segmented regression indicated no detectable change in establishment rate patterns during the Euro-American settlement period (approx. 1860-1900 CE), with a continuation of the upward trend that began in the late 1700s CE. The largest recent change in establishment rates occurred following ∼1920 CE, when both the 20-year average and segmented regression indicated establishment rates began to decline (Table S2). Surprisingly, we found that by the late 20^th^ century tree establishment rates had reached their lowest level in 400 years, 0.16 (0.15-0.17 95% CI) per 20-year interval.

To better quantify the mechanisms explaining recent increases in total establishment, we separated contributions resulting from steady multiplicative population growth alone, from contributions of changing establishment rates. We did so by comparing two simulations for each of the 29 populations: one used the observed establishment rates for each population from 1600-1980 CE (i.e. “changing establishment rates”), while the other was a null model that maintained a constant establishment rate using the average establishment rate for each population from 1600-1980 CE (i.e. “constant establishment rate”). We found that multiplicative population growth with a constant establishment rate alone consistently predicted most of the observed establishment increases across decades at a regional scale (Fig 3A,B). Hence, increased establishment rates due to anthropogenic or other environmental drivers are not needed to explain observed increases in establishment and density beginning in the 1800’s CE. Multiplicative population growth alone did not always predict observed temporal variability in establishment in an individual population, indicating that anthropogenic or other environmental drivers may play an important role in dictating establishment at a local scale. However, these positive and negative deviations were found throughout the 400-year period, rather than being concentrated only during the late 1800’s CE (Fig. 3C). For example, constant establishment rates underpredicted establishment amounts from ∼1600-1700 CE by up to 80% on average while only underpredicting establishment by ∼25% from 1900-1920 CE.

**Figure 3.**
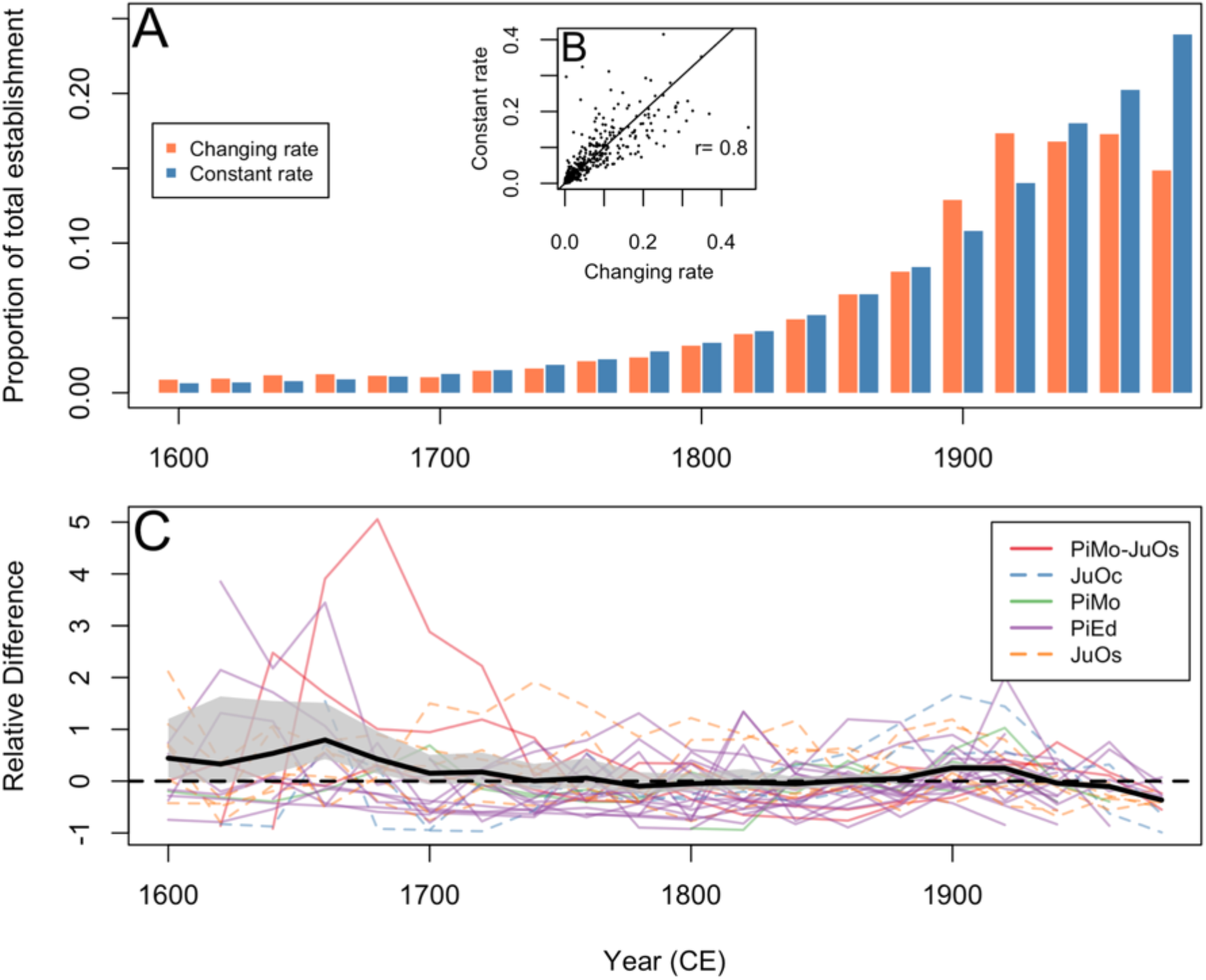
Predictability of tree establishment patterns since 1600 CE. (A) Bars representing average expected establishment in 20-year intervals across all populations expected from changing per-capita establishment rates (orange) and multiplicative population growth with a constant establishment rate (blue). (B) Expected establishment under changing and constant establishment rates across all populations and correlation of the establishment from each. Points represent posterior median estimates for year population each 20-year interval after 1600 CE. (C) Relative difference 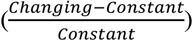 in establishment between each mechanism for each population. Solid black line indicates average and 95% CI. Values of 0 (dashed black line) indicate that the amount of establishment is the same under either mechanism, whereas values above 0 indicate that changing establishment rates would be expected to lead to greater establishment.

### Robustness of establishment rate estimates

Tree ring derived age structure data can suffer from a “fading record” problem, where data reliability, and thus ecological inference, declines further in the past^16^. Trees that were previously present in a population may have died before being observed, and probability of missing trees is likely to increase as we move further back in the historic record. As a result, tree survival rates could have impacted our inference of past establishment rates. Our Bayesian inference approach allowed us to test this and account for uncertainties in survival rates in estimates of establishment rates. We found that despite high levels of uncertainty in estimates of past survival rates, establishment rate estimates were typically well constrained and accurately identifiable (Figs. 4, S3, S4). While uncertainty in establishment rates increased with time since present (Figs. 4B, S3B), estimates of survival rates derived from posterior distribution samples varied independently from establishment rates, showing that uncertainties about survival rates had little impact on our estimates of establishment rates (Fig. 4C). Deterministic population models help illustrate why this occurred. Establishment rate estimates remained similar in the face of disparate survival scenarios because, when mortality occurred, observed age structures underestimated both past establishment (i.e. many of the established plants were not still present to be censused) and past population size, which had a cancelling effect when calculating per-capita establishment (Fig. S5). Similarly, reproductive lags in young trees did not change overall establishment rate results (Fig. S6).

**Figure 4.**
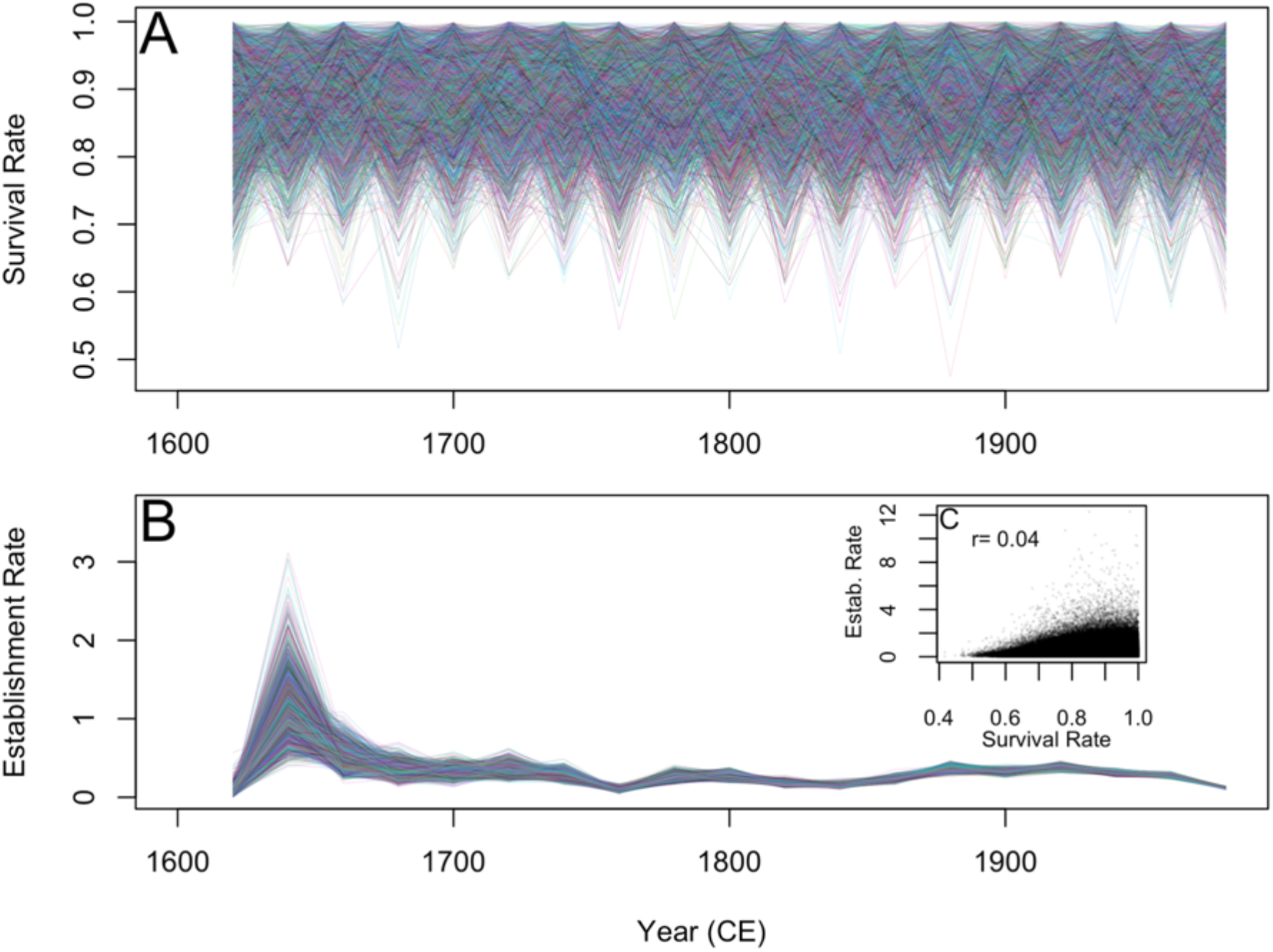
Impacts of survival rates on estimated per-capita establishment rates. (A) Lines representing individual posterior draws for survival rates for a single site illustrating wide variability. (B) Lines representing individual posterior draws for establishment rates for the same site illustrating well constrained estimates. (C) Correlation of survival and establishment rate posterior draws in all sites illustrating low correlation.

### Long-term drivers of woody plant abundance

We found little evidence to support the hypothesis that an unprecedented increase in tree establishment rates drove observed regional increases in tree density in the 19^th^ and 20^th^ century. Woodland density and extent may have actually been steadily increasing for at least the last 400 years. Such sustained population growth may have been driven by late Holocene conditions, as shown by paleoecological records from the western USA^24–26^. The period from 900 to 1500 CE was associated with region-wide increases in drought and wildfire activity and with declining presence of forests and woodlands, the extent of which may exceed any conditions in the modern era^8,26–28^. Paired archaeological and paleoecological data from the Great Basin and Colorado Plateau region also indicate that fuelwood harvesting, agriculture, and cultural fire practices may have led to further declines in pinyon-juniper abundance and extent prior to 1500 CE^24,29,30^. Woodland harvesting and contraction prior to 1500 CE would have created a large unfilled niche space for pinyon-juniper woodlands, ideal for sustained range-wide tree population increases and range expansion once climate conditions became cooler and wetter and cultural practices changed (Fig. S2B; *32*). This hypothesis is further supported by pollen and midden data from the western USA which indicate tree abundance began to increase well before the 19^th^ century as well as evidence of general range expansion of pinyon and other dryland tree species throughout the last 500-1000 years^6,28,31,32^.

Pinyon-juniper woodlands have undergone several periods of growth, expansion, and migration throughout the Holocene, often punctuated by periods of contraction^6,25,28,33^. The period of range expansion since 1600 CE may simply represent the latest period in this range expansion-contraction dynamic. Wildfire continues to play an important role in structuring pinyon-juniper woodlands. However, unlike other ecosystems known for frequent, low-severity fires, fire regimes in pinyon-juniper woodlands have generally been characterized by infrequent, high-severity fires (fire rotations of 400+ years), and several of the sites included in our study from the Colorado Plateau region show no evidence of fires in the last 400 years^34,35^. As a result, recent fire exclusion in pinyon-juniper woodlands is unlikely to explain accelerating increases in tree density^3^.

We found that the dominant trend since 1900 CE is declining, not increasing, tree establishment rates, and that establishment rates are currently at their lowest level in the past 400 years. The 1930’s CE were a period of drought and anomalously warm conditions throughout much of the western US, and have been subsequently followed by variable, but generally warming temperatures and extreme drought attributed to anthropogenic climate change^27^. Previous studies from higher elevation forests in the western USA have found declines in tree survival across the 20^th^ century and hypothesized that climate change was a likely cause. Our results and other contemporary studies in pinyon-juniper woodlands^36,37^ suggest that climate change may be a factor contributing to the unprecedented decline in tree establishment rates we observed. Declining establishment rates could also simply represent density-dependence as stands reach their carrying capacity, although declining establishment rates have been observed across a range of stand densities and ages^8^. Regardless of the underlying mechanism, finding that woodland establishment rates are at their lowest rate in at least 400 years provides important, and previously unrecognized, context for understanding the dynamics of woodland ecosystems.

Increases in woody plant density and extent have been noted in many dryland ecosystems across the world^2^. Our results suggest that intrinsic population processes alone can explain the rapid change of dryland ecosystems from low, but seemingly stable levels of woody plant density, to increasingly dense woody vegetation dominated primarily by young plants. Age structure data on dryland woody plants over the last several hundred years are rare outside of the southwest USA but similar mechanisms may be operating in other regions. Estimated age structures from a savanna site in southern Texas, USA^40^, show rapid increases in tree density in the 20^th^ century despite per-capita establishment rates that fall well within the range of variability from 1770-1900 CE (Fig. S7), suggesting multiplicative population growth could explain rapid increases in woody plant density. Fossil pollen data from Ethiopia and the Chihuahuan desert indicate declines in woodland abundance during the medieval climate anomaly (∼950-1250 CE), followed by the recovery of woody vegetation beginning during the little ice age (∼1400-1700 CE) ^41,42^. Research from African savannas also suggest that historic conditions, and not the current environment, may be the primary driver of extant woody plant abundance^43^.

Our findings have important implications for the management and restoration of pinyon-juniper woodlands, one of the most widespread vegetation types in the USA. Removing pinyon and juniper trees and returning historic ecosystem services provided by grasslands and shrublands has become an increasingly high priority for land management agencies, often based on the assumption that increases woodland density and extent over the last 200 years represent an unnatural process^21,44^. Our results suggest that the processes driving recent increases in woodland density may not be historically unprecedented, but rather a continuation of population recovery and expansion that initiated at least 400 years ago, possibly following major changes in western North American forests caused by drought, wildfire and shifts in Indigenous land-use^26,27,29^. As a result, management practices intending to address woody encroachment by removing trees at local scales cannot address the underlying regional drivers of encroachment in these ecosystems. Despite this, widespread increases in woody plant density may still alter the provision of important ecosystem services or habitats for species of concern^19^. Thus, it is crucial to consider the underlying processes driving increasing woody plant density, the consequences of greater woody plant density for ecosystem processes and services, as well as projected future conditions given likely scenarios of climate and disturbance. In fact, management should carefully consider the possibility of future woodland contraction. Given that pinyon-juniper establishment rates are nearing their lowest level in 400 years and that widespread mortality has occurred in the Southwest US, we may be entering a period when population growth can no longer be maintained, and range contraction may be more likely^37,45^.

Our research also provides a cautionary tale for ecologists studying how anthropogenic factors influence ecological processes, particularly those associated with Euro-American settlement in the western US. Human activities have undoubtedly changed the structure and function of ecosystems, including the introduction and spread of invasive species and the widespread removal of native vegetation^46,47^. At the same time, impacts of settlement occur against the backdrop of long-term changes in climate and historical legacies that are critical to understanding the modern distribution and abundance of species, but are often not readily apparent today^16,25^. As a result, recent observed shifts in woody plant density and cover may coincide with changing environmental conditions during the settlement period, but may not be causally linked. Disentangling the causes of historical vegetation change can be challenging, and it can lead ecologists to seek explanations that invoke complex interactions among global change agents. Our study highlights the need to compare such hypothesized models of change with simpler and often more parsimonious explanations, including null models of population dynamics in the absence of external drivers such as climatic changes, wildfire regime shifts, or land-use changes.

## Materials and Methods

### Demographic background

Our methods are based on classical population ecology theory, so it is helpful to begin with a little background. Changes in population size over time can be described by

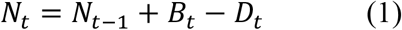

Where *N*_*t*_ is the population size at time *t, B*_*t*_ is the number of individuals establishing in the population and *D*_*t*_ is the number of deaths. But this simple formulation fails to capture the compounding, multiplicative nature of population growth. Each new individual established will eventually have an opportunity to reproduce if they survive, further expanding the population and increasing establishment. So, it is more useful to express population change in per-capita establishment (*b*_*t*_) and survival rates (*s*_*t*_):

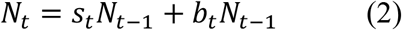

The number of trees of a given age (*B*_*t*_, i.e. each bar in Fig. 2A) represents *b*_*t*_*N*_*t*-1_, the product of both the per-capita establishment rate and the population size. Thus, we can observe greater establishment by either having a larger population size or increasing the average number of offspring per individual. As long as establishment rate (*b*_*t*_) exceeds the mortality rate (1 − *s*_*t*_) the number of establishing individuals is expected to increase over time simply because each successive cohort contributes more reproductive potential, even if *s*_*t*_ and *b*_*t*_ are constant. Because population size itself influences the total amount of establishment, demographers look at changes in per-capita establishment (*b*_*t*_), not total establishment (*B*_*t*_) to identify the impacts of changing conditions such as changes in disturbance, climatic conditions, and other abiotic and biotic drivers ^12,13^. It’s worth noting that authors of previous pinyon-juniper studies often refer to *b*_*t*_*N*_*t*-1_ as the establishment rate^8,48^, but in those cases, this is the number of new trees per unit time, not the demographic rate (new trees per existing tree per unit time). If changes in climate, disturbance, or other abiotic and biotic conditions were driving the acceleration of establishment (*B*_*t*_), this would be reflected by an increase in the per-capita establishment rate (*b*_*t*_). Rearranging equation 2, the per-capita establishment rate for any period can be empirically calculated as:

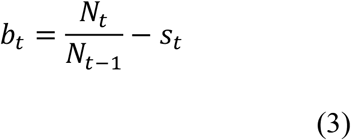

### Age Structure Data

To understand how per-capita establishment rates have changed over time we digitized and collated age structure data from existing studies. Studies included those previously known to the authors and additional published articles, reports, and theses identified as including age structure data using Google Scholar and Web of Science searches with combinations of the phrases “pinyon”, “piñon”, “juniper”, “age structure”, and “establishment”. Papers that only sampled large trees, or in other ways incorporated a biased sampling of age structure, were not used. Because we extracted data from graphs, we could not use papers where graphs were impossible to interpret and extract data with confidence. In addition, any study that aggregated tree establishment dates into intervals greater than 20 years was not included because of lack of temporal resolution. Because we are interested in how establishment rates have changed prior to and after Euro-American settlement, we did not include suitable age structures from studies where all trees established after 1800. This resulted in datasets from 29 populations at 23 sites (some sites have more than one species measured) from 10 separate studies (Table S2). Data points were extracted from published figures using Web Plot Digitizer ^49^, an online application for extracting data from scientific figures. Each extracted point included a very small amount of numerical error associated with digitization. Because in each dataset establishment years were aggregated into 10- or 20-year bins, we were able to correct numerical digitization errors in years of establishment. The numerical errors in establishment amount data were small (Fig. S8). Each study calculated tree ages within stand plots or transects using standard dendrochronology approaches and aggregated trees into 10-or 20-year cohorts. 12 of the datasets did not age the smallest trees in the stand (e.g. no trees smaller than 3 cm diameter were aged); in these datasets we did not use any tree establishment data after the 1940-1960 interval so as to not artificially lower establishment rates because of missing trees. Two datasets did not include trees < 4cm diameter, but already excluded data in their reporting after 1950 to account for missing small trees ^10^. Datasets spanned a diversity of geographic locations in the Great Basin and Colorado Plateau (Arizona, California, Colorado, Idaho, Nevada, and Utah; Fig. S1) and species including *Pinus monophylla* (PiMo), *Pinus edulis* (PiEd), *Juniperus osteosperma* (JuOs), *Juniperus occidentalis (*JuOc), and mixed stands dominated by *Pinus monophylla*, with a minor *Juniperus osteosperma* component (PiMo-JuOs).

To directly compare all datasets at the same time scale we aggregated 10-year datasets to 20-years by summing establishment in two 10-year intervals. In some cases, 10-year datasets did not end on the full 20-year interval, and only quantified establishment during the first 10 years (e.g. 1960-1969) of the 20-year interval (1960-1979). In these cases, the last 10 years of the dataset were not included in the analysis avoid underestimates of establishment (e.g. using 10 years of data for a 20-year interval). In cases where authors reported relative establishment (e.g. proportion of trees in each age cohort, trees in a standardized area), establishment values were converted back to tree counts using reported sample sizes or search areas. This step was necessary for our Bayesian inference approach where uncertainty depends, in part, on the number of individuals sampled.

### Establishment rate inference

We used a Bayesian inference approach to estimate establishment rates in each population. This approach allowed us to account for full uncertainty in unknown survival rates and population size (*s*_*t*_, *N*_*t*_) when estimating establishment rates (*b*_*t*_).

Population size at each time step was determined by eq. 2

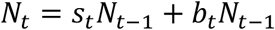

where *N*_0_, the number of founder trees before first recorded tree, was treated as an unknown parameter with prior *Normal*(50,10).

The underlying population dynamics model (eq. 2) was then connected to observed establishment data in each population as

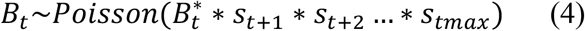

Where *B*_*t*_ is the observed number of individuals that established at time *t*, 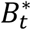 is the true number of individuals that established (*b*_*t*_*N*_*t*-1_) corrected for the cumulative survival of that cohort from establishment to observation. Given observations of average decadal survival rates from 95% to >99% in pinyon-juniper species^50^, we used a truncated normal informative prior for all intervals and populations of *s*∼*Normal*(0.95,0.1)*T*[0,1]. This prior still allowed for considerable uncertainty in survival rates that aligns with current observed variability ranging from 50% to near 100% survival (e.g. Fig. 4). Establishment rates were given non-informative uniform priors. Models were fit using “rstan” ^51^ and model convergence was checked visually and using r-hat statistics (r-hat <1.1).

To test the ability of our modeling approach to recover underlying establishment rates, we simulated 100 establishment datasets over 30-time intervals using randomly generated, known survival and establishment rate values, and then fit the model described above to test if posterior estimates of establishment rates align with the true values. The ability of our model to recover true establishment rates was calculated by using the correlation between true establishment rate values and estimated posterior median establishment rate values, root mean squared error (RMSE) between true and estimated establishment rate values across time periods (lower values indicate higher accuracy), and coverage. Coverage calculates the proportion of the true establishment rate values that fall within credible intervals, with the expectation that a well developed, accurate model should recover the true value in its 95% credible interval approximately 95% of the time. Results indicated that while uncertainty in establishment rates increased with time since present, true establishment rates were identifiable, with a correlation of 0.78 between posterior median establishment rates and true values (Fig. S3A). Across simulated datasets RMSE was highest earliest in the timeseries (1.31), but declined rapidly nearing the present (0.03) (Fig. S3B). Even when establishment rate estimates were less precise, true values were still accurately contained within posterior distributions as indicated by coverage values near 0.95 throughout the timeseries (Fig. S3C).

We calculated the mean per-capita establishment rate for each 20-year interval. We also performed segmented regression to identify and quantify major changes in per-capita establishment rates over time using the “segmented” package in R ^52^. Segmented regression identifies the location and slope of change points in a response variable given a fixed number of change points. We performed segmented regression between the estimated establishment rates in all plots (response variable) and interval year (explanatory variable) with one to four change points and calculated AIC scores to determine the best model.

### Deterministic establishment rate calculations

To complement our Bayesian inference approach and better understand observed demographic results, we also calculated establishment rates deterministically using equation 3. This approach required us to assume survival rates for each population. To do this we tested three survival scenarios: constant 100% survival, constant 90% survival, and simulated varying survival rates. 90% survival over each 20-year interval is comparable to the current survival rates of *Pinus monophylla* and *Pinus edulis* ^37^, while juniper species typically have higher survival. Simulated survival rate time series were generated using a forward and reverse random walk starting at logit(0.95), that was then inverse-logit transformed to the 0-1 scale.

We corrected observed establishment numbers for past survival by accounting for the cumulative survival cohort from when the establishment occurred in interval, *t*, to the time of observation (*tmax*) as:

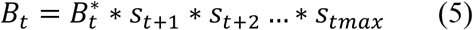

Here *B*_*t*_ is again the observed number of established individuals in interval, *t*. 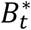 is the true number of individuals that established in that interval.

This can be rearranged to calculate the true number of individuals and simplified as

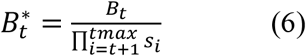

In other words, the true number of individuals that established in a past year,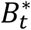, is the observed number of establishing trees at that time corrected for the cumulative survival rate of that cohort from the time of establishment to the observation. Finally, we calculated the true population size at any time 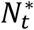 as

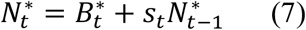

The deterministic and Bayesian approach yielded similar estimates of establishment rates (correlation of 0.78), however because the deterministic approach did not allow for uncertainty it often results in more extreme values (Fig. S9).

### Delayed reproduction

Time to first reproduction for pinyon and juniper can vary by species and site, but can take multiple decades. This could impact inferred per-capita establishment rates because not all members of the population would be contributing to new tree establishment. To understand how delayed reproduction would have impacted our inferred per-capita establishment rates we deterministically calculated per-capita establishment rates assuming a 40-year lag (two time intervals) in reproduction. To do this we created a simple structured population model where we separated the population into those contributing to reproduction and those contributing to survival.

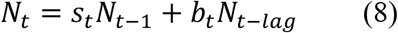

Here *N*_*t*_ is the population size at interval, *t*, and *N*_*t* − *lag*_ is the population size at an interval in the past corresponding to the lag. This can be rearranged as

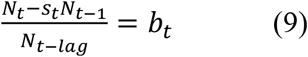

Assuming the survival rate is equal to 1, this simplifies to

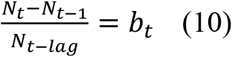

*N*_*t*_ − *N*_t-1_ is equal to the number of newly established trees observed in an interval, *B*_*t*_, thus the per-capita establishment rate is

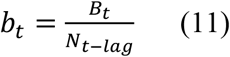

In other words, the per-capita establishment rate is the number of newly established trees at *t* divided by the population size at *t* − *lag*.

### Decomposing contributions to establishment

To quantify the role of multiplicative population growth relative to changing per-capita establishment rates in driving age structures, we simulated expected establishment in each 20-year interval from 1600-1980. We quantified the role of multiplicative growth by simulating population growth under a constant establishment rate, using the mean of the observed establishment rate from 1600-1980 for that population. This quantified the amount of establishment we would expect if no change in establishment rates occurred and multiplicative population growth alone was responsible for observed dynamics. We then repeated this simulation, but with varying per-capita establishment rates, using the observed establishment rate for each 20-year interval from each population. This quantified the amount of establishment we would expect with both multiplicative growth and changing per-capita establishment rates. Similar to Fig. 2A, expected observed establishment was then calculated as the survivorship of each 20-year cohort to observation and relativizing to the total for each population and simulation to allow for direct comparison across populations. The difference in establishment between these two simulations is the contribution of changing per-capita establishment rates to overall establishment. In all cases, simulations accounted for and incorporated survival rate and establishment rate uncertainty by iterating over the draws from the posterior distribution for each population.

## Acknowledgments

We thank the authors that generated the data used in this work, which made this research possible. We thank Peter Adler for helpful feedback on earlier versions of this manuscript. This research was funded by the USDA National Institute of Food and Agriculture, the Nevada Agricultural Experiment Station, and by the USDA Forest Service, Rocky Mountain Research Station. The findings and conclusions in this publication are those of the authors and should not be considered to represent any official USDA determination or policy.

## Author Contributions

RKS: Conceptualization, Data Curation, Formal Analysis, Writing. EP: Data Curation, Formal Analysis, Writing. FB, AU, PJW: Writing.

## Data availability

Data are available on GitHub at https://github.com/bobshriver/PJStructure.

Data will be permanently archived on Zenodo upon acceptance.

## Code availability

R code to reproduce all analyses are available on GitHub at https://github.com/bobshriver/PJStructure. Code will be permanently archived on Zenodo upon acceptance

