## Supplemental Material for "Long-term tree population growth can predict woody encroachment patterns"

### Supplement

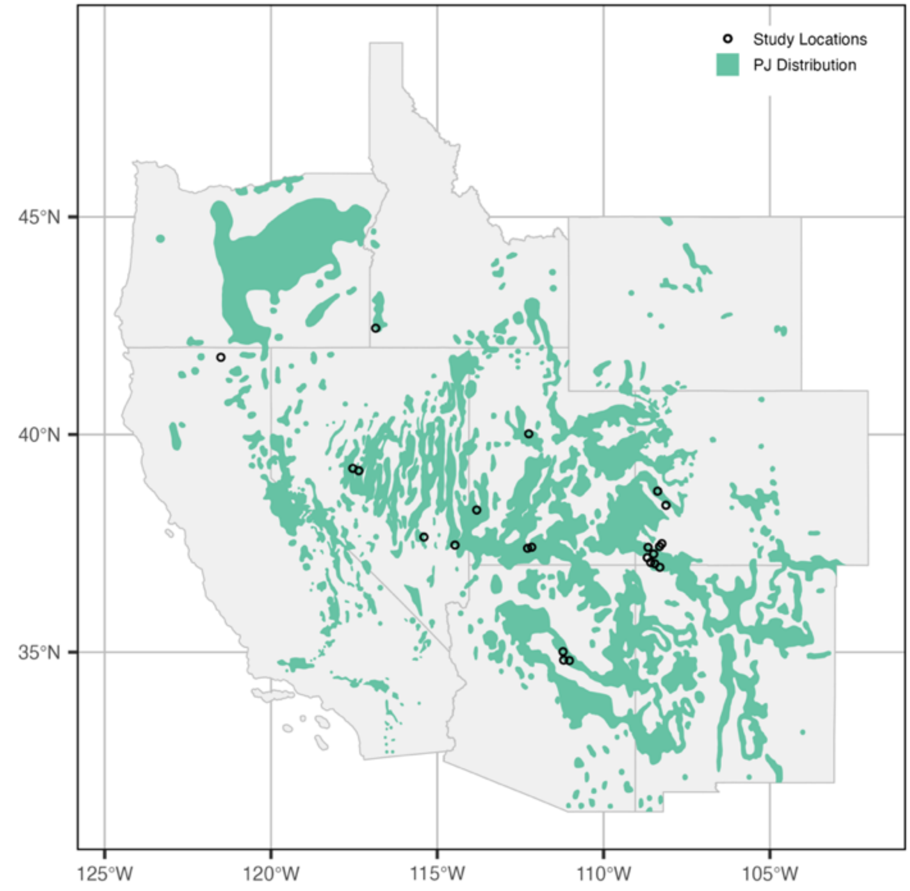


**Figure S1. Map of study locations.** Points indicate individual study sites. Nearby, overlapping site points were jittered to make each point visible.

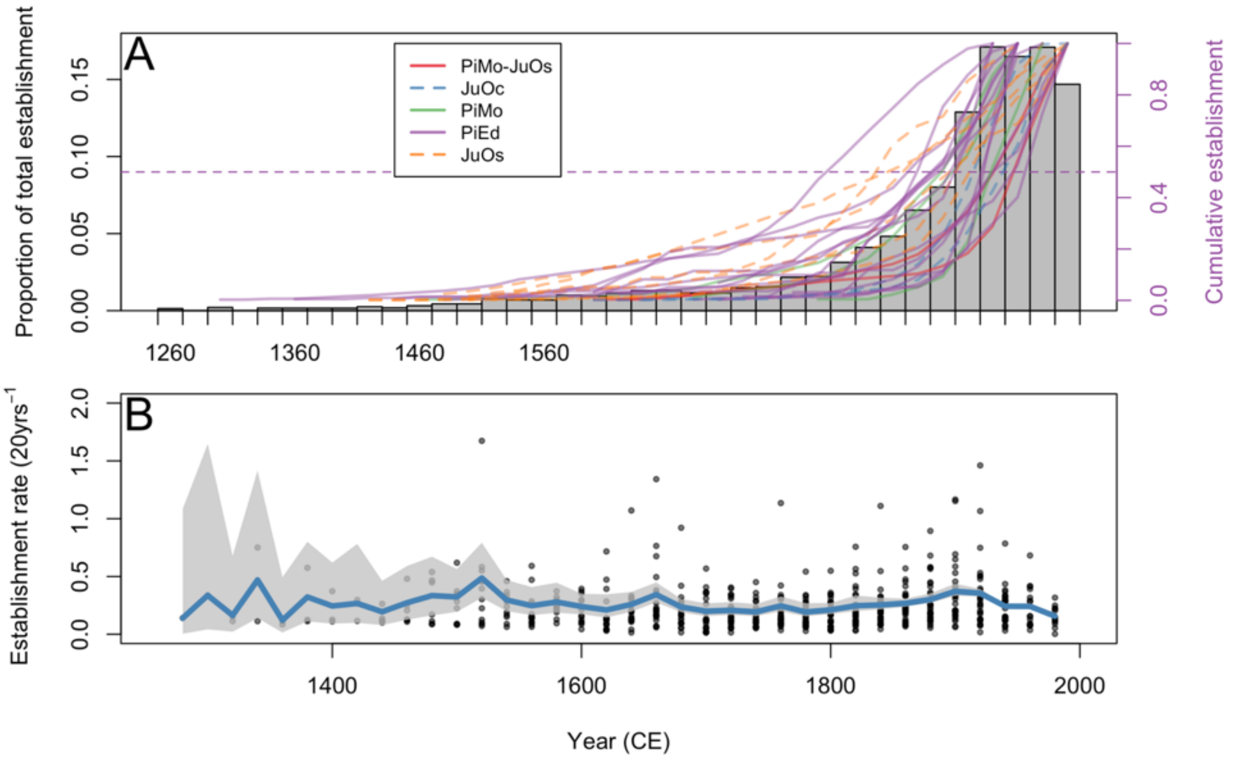


**Figure S2. Tree establishment over entire timeseries.** (A) Proportion of total establishment occurring in each 20-year interval averaged across 29 populations and cumulative establishment in each of the 29 populations broken down by species. (B) Per-capita establishment rates for each population (points), 20-year average (blue) and 95% CI (grey).

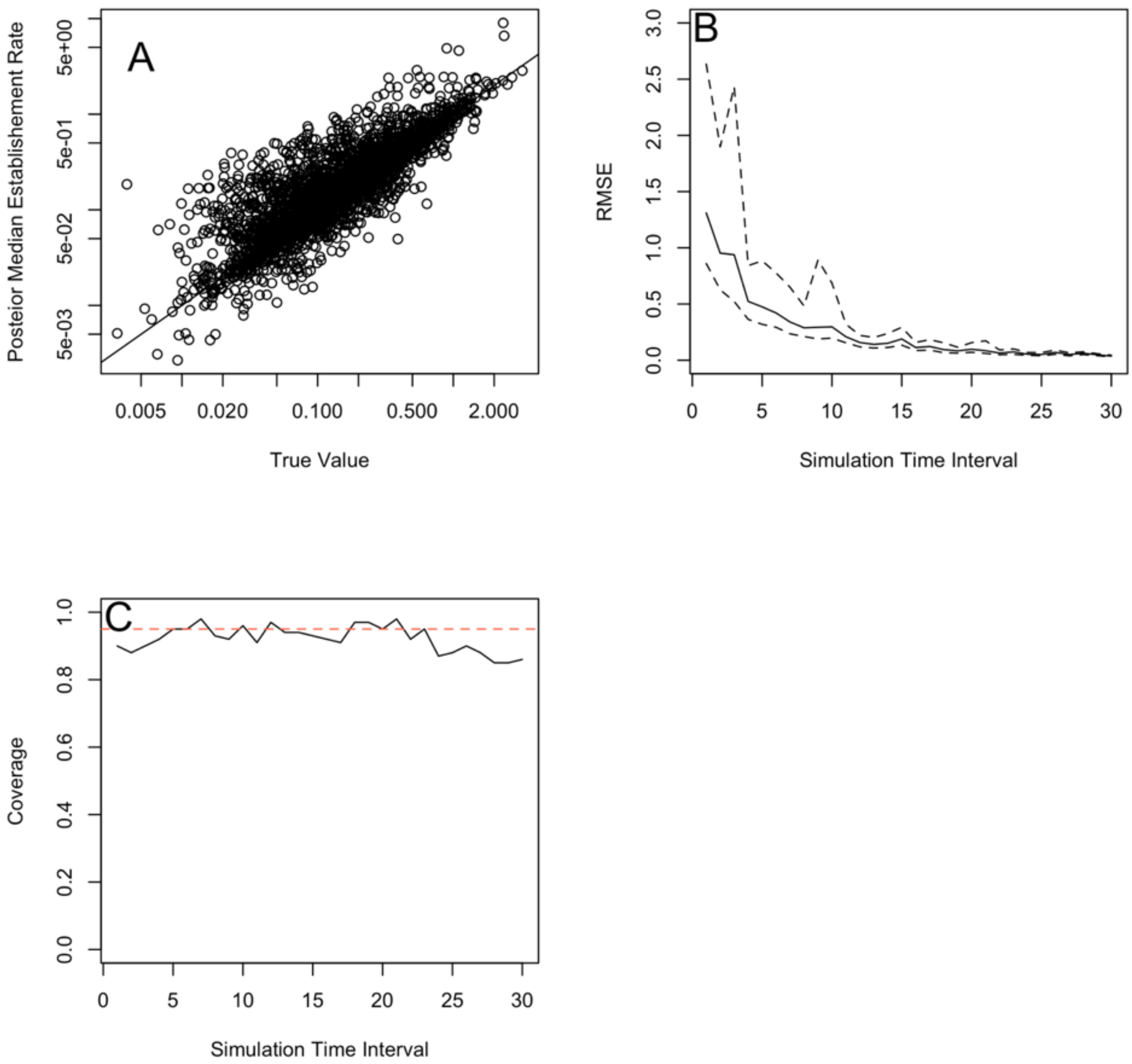


**Figure S3. Ability of Bayesian model to estimate establishment rates.** (A) Scatter plot showing the true establishment rates and posterior median estimates for each of the 100 simulated datasets across all 30-time intervals. Black line is the 1:1 line. (B) Root mean squared error (RMSE) between the estimated and true establishment rate across time intervals. Solid line is posterior median and dashed lines are 95% CI. 0 interval on x-axis represents the furthest past and 30 the most recent. (C) Coverage of true establishment rate values in posterior estimates. The dashed red line is the target, 0.95. The solid black line is the estimated coverage.

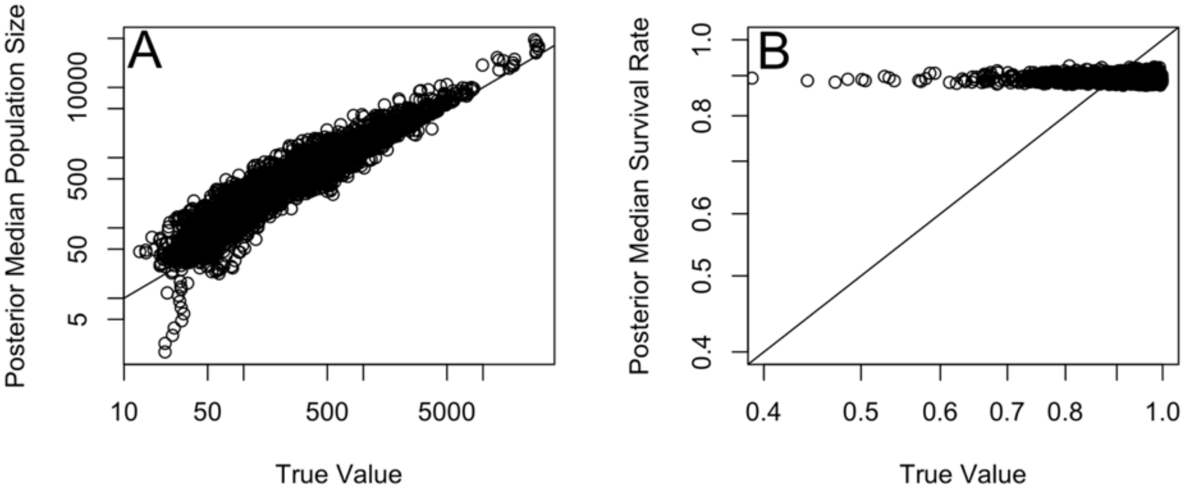


**Figure S4. Ability of Bayesian model to estimate population size and survival rates.** (A) Scatter plot showing the true population size and posterior median estimates for each of the 100 simulated datasets across all 30-time intervals. Black line is the 1:1 line. (B) Scatter plot showing the true survival rates and posterior median estimates for each of the 100 simulated datasets across all 30-time intervals. Black line is the 1:1 line.

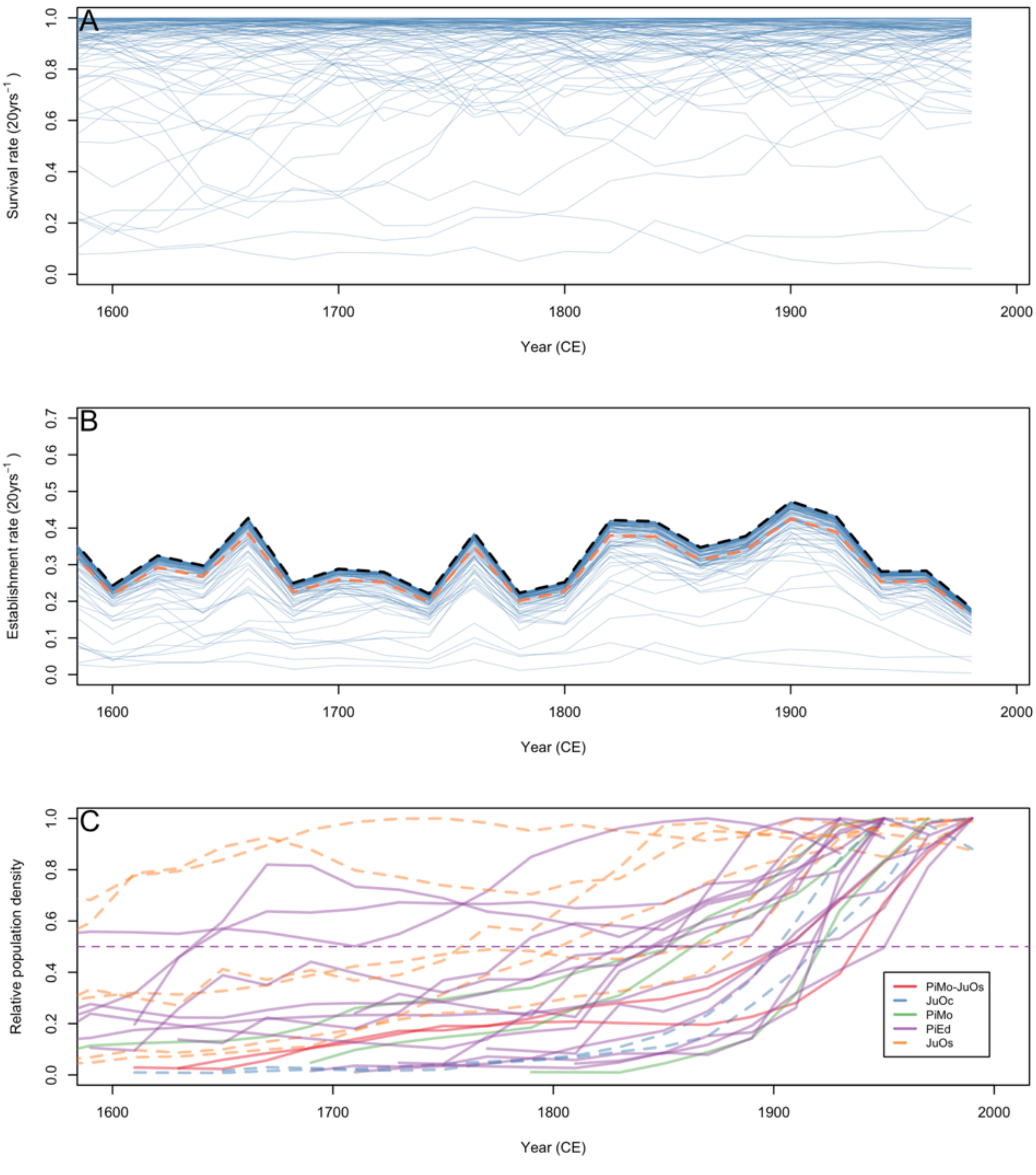


**Figure S5. Impacts of survival rates on estimated per-capita establishment rates.** (A) 100 randomly generated survival timeseries. (B) Estimated per-capita establishment rates given each of the survival scenarios in panel A. Dashed black and orange line indicates establishment rates with constant 100% and 90% survival, respectively. (C) Population density in each of the 29 populations broken down by species assuming 90% survival, illustrating increases in past population sizes with lower survival relative to 100% survival (Fig 2A). Dashed line indicates 50% of total.

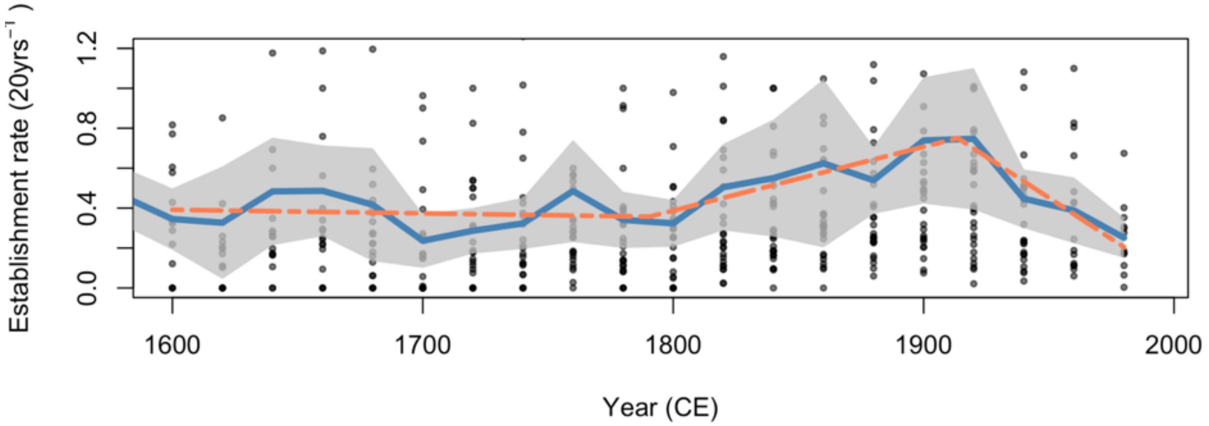


**Figure S6. Establishment rates given a 40-year lag in reproduction for new trees.** Per-capita establishment rates for each population (points), 20-year average (blue) and 95% CI (grey), and results of segments regression (dashed orange).

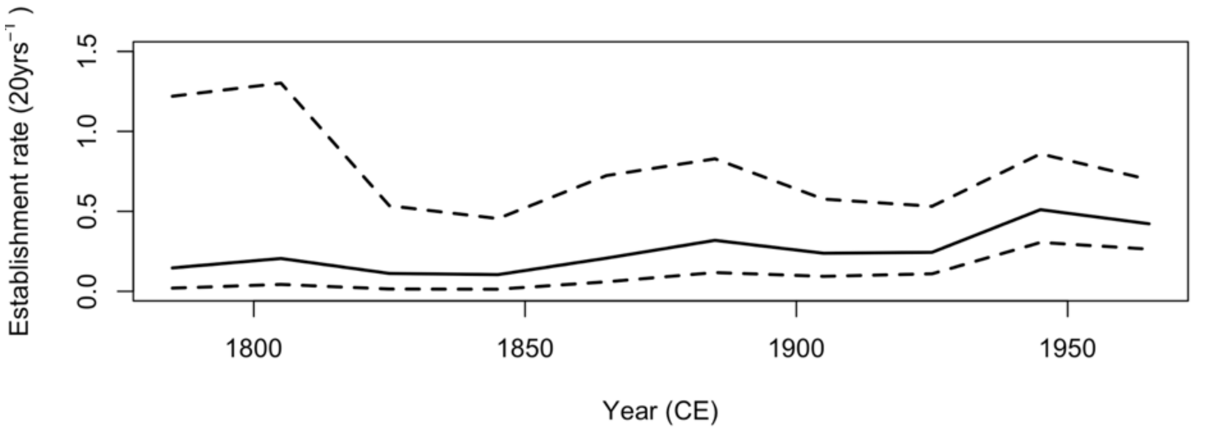


**Figure S7. Establishment rates in mesquite savanna of Texas, US.** Solid line indicates posterior median estimate. Dashed lines are 95% credible intervals.

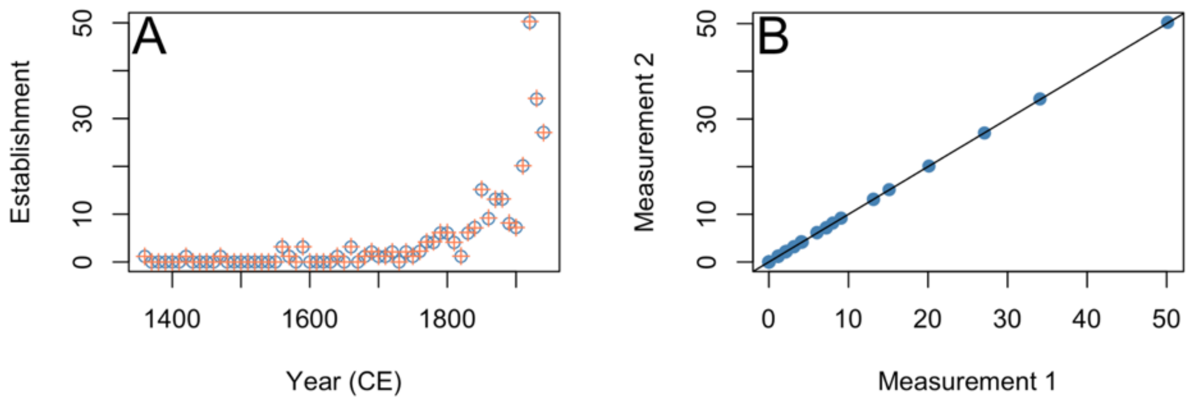


**Figure S8. Example of errors associated with plot digitization.** A) Comparison of two measurements over time. One measurement is indicated with the orange cross and the other the blue circles. B) Comparison of measurements with 1:1 line indicating an exact match.

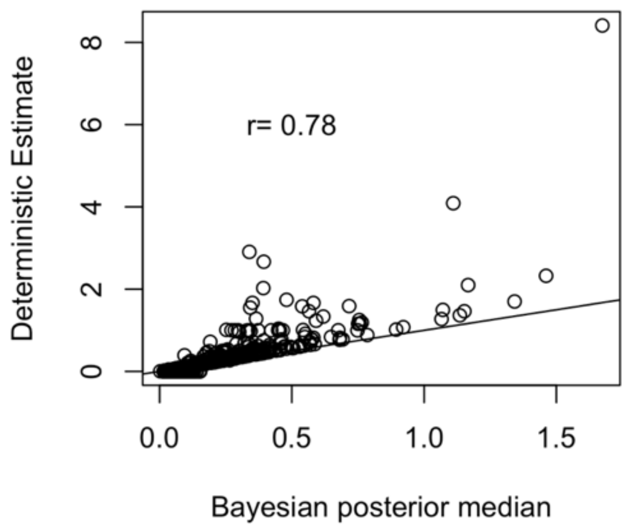


**Figure S9. Correspondence between Bayesian and deterministic survival rate calculations.** Points represent individual 20-year intervals in each population. Solid line is the 1:1 line.

**Table S1. Description of datasets used, and reason for omission for those not used.**

| Paper | Figure | Used | If not used, reasoning | # of Sites | Species |
|---|---|---|---|---|---|
| Miller, R. F., R. J. Tausch, E. D. McArthur, D. D. Johnson, and S. C. Sanderson. 2008. Age structure and expansion of pinon-juniper woodlands: a regional perspective in the Intermountain West. Page RMRS-RP-69. U.S. Department of Agriculture, Forest Service, Rocky Mountain Research Station, Ft. Collins, CO. | Fig. 4 | YES | Only mixed age stands used, with trees before 1800. | 3 | 1)PiMo-JuOs, 2)JuOc |
| Biondi, F., and M. Bradley. 2013. Long-term survivorship of single-needle pinyon (Pinus monophylla) in mixed-conifer ecosystems of the Great Basin, USA. Ecosphere 4:art120. | Fig. 4 & Fig. 5 | YES | 0.1 ha plots | 2 | 1)PiMo, 2)JuOs |
| Barger, N. N., H. D. Adams, C. Woodhouse, J. C. Neff, and G. P. Asner. 2009. Influence of Livestock Grazing and Climate on Pinyon Pine (Pinus edulis) Dynamics. Rangeland Ecology & Management 62:531–539. | Fig. 1 | YES | | 2 | 1)PiEd |
| Bauer, J. M. 2006. Fire history and stand structure of a central Nevada pinyon-juniper woodland. M.S., University of Nevada, Reno, United States -- Nevada. | Fig. 8 | YES | | 1 | 1)PiMo-JuOs |
| Shinneman, D. J., and W. L. Baker. 2009. Historical fire and multidecadal drought as context for piñon–juniper woodland restoration in western Colorado. Ecological Applications 19:1231–1245. | Fig. 5 | YES | | 1 | 1)PiEd, 2)JuOs |
| Landis, A., and J. D. Bailey. 2005. Reconstruction of age structure and spatial arrangement of piñon–juniper woodlands and savannas of Anderson Mesa, northern Arizona. Forest Ecology and Management 204:221–236. | Fig. 3 | YES | | 3 | 1)PiEd, 2)JuOs |
| Eisenhart, K. S. 2004. Historic range of variability and stand development in piñon-juniper woodlands of western Colorado. Ph.D., University of Colorado at Boulder, United States -- Colorado. | Fig. 3.5 | YES | Only Dominguez WSA used, which sampled all pines. Rest of sites only sample subset of large trees. | 1 | 1)PiEd |
| Tausch, R. J., and N. E. West. 1988. Differential Establishment of Pinyon and Juniper Following Fire. The American Midland Naturalist 119:174–184. | Fig. 2 | YES | Note: Numbers were exp transformed and differenced to get establishment | 1 | 1)PiMo, 2)JuOs |

| | | | | | |
|---|---|---|---|---|---|
| | | | numbers in each intervals. Intervals offset by 5 years, so aligned with 5 year mismatch. | | |
| Floyd, M. L., D. D. Hanna, and W. H. Romme. 2004. Historical and recent fire regimes in Piñon–Juniper woodlands on Mesa Verde, Colorado, USA. Forest Ecology and Management 198:269–289. | Fig. 3 | YES | | 8 | 1)PiEd |
| Miller, R.F., E.K. Heyerdahl, and K. Hopkins. 2003. Fire regimes, pre- and post-settlement vegetation, and the modern expansion of western juniper at Lava Beds National Monument, California. Final Report to the USDI Lava Beds National Monument. | Fig 3. | YES | | 1 | 1)JuOc |
| Miller, R. F., and J. A. Rose. 1995. Historic expansion of Juniperus occidentalis (western juniper) in southeastern Oregon. GREAT BASIN NATURALIST 55. | Fig 1. | NO | Line plots don't differentiate observed data points from point interpolation | NA | NA |
| Blackburn, W. H., and P. T. Tueller. 1970. Pinyon and Juniper Invasion in Black Sagebrush Communities in East-Central Nevada. Ecology 51:841–848. | Fig. 6 | NO | Line plots don't differentiate observed data points from point interpolation. Most post-1800 | NA | NA |
| Waichler, W. S., R. F. Miller, and P. S. Doescher. 2001. Community Characteristics of Old-Growth Western Juniper Woodlands. Journal of Range Management 54:518–527. | Fig 2. | NO | We limited the re-analysis to 10- and 20-year intervals, 50-year intervals in this analysis. | NA | NA |
| Tonnesen, A. S., and J. J. Ebersole. 1997. Human Trampling Effects on Regeneration and Age Structures of Pinus Edulis and Juniperus Monosperma. The Great Basin Naturalist 57:50–56. | Fig. 1 | NO | In heavily used park. Recent and continued trampling of seedlings and saplings by park users. | NA | NA |
| Kitchen, S. G. 2012. Historical fire regime and forest variability on two eastern Great Basin fire-sheds (USA). Forest Ecology and Management 285:53–66. | Fig. 6 | NO | sampling methods were restricted to >20cm DBH | NA | NA |

| | | | | | |
|---|---|---|---|---|---|
| Floyd, M. L., W. H. Romme, D. P. Hanna, and D. D. Hanna. 2017. Historical and Modern Fire Regimes in Piñon-Juniper Woodlands, Dinosaur National Monument, United States. Rangeland Ecology & Management 70:348–355. | Fig. 2 | NO | We limited the re-analysis to 10- and 20-year intervals, 25-year intervals in this analysis. | NA | NA |
| Margolis, E. Q. 2014. Fire regime shift linked to increased forest density in a piñon–juniper savanna landscape. International Journal of Wildland Fire 23:234–245. | Fig. 4 | NO | Only largest trees were included in analysis. | NA | NA |
| Soulé, P. T., P. A. Knapp, and H. D. Grissino-Mayer. 2004. Human Agency, Environmental Drivers, and Western Juniper Establishment During the Late Holocene. Ecological Applications 14:96–112. | Fig. 7 | NO | Unable to differentiate between individual year bars. | NA | NA |
| Swetnam, T. W., and J. L. Betancourt . 1998. Mesoscale disturbance and ecological response to decadal climatic variability in the American Southwest. *Journal of Climatology* **11**: 3128–3147. | Fig. 15 and 17 | NO | Fig. 15 only old trees. Fig. 17 was only post 1900 | | |
| Andre, G. St., H. A. Mooney, and R. D. Wright. 1965. The Pinyon Woodland Zone in the White Mountains of California. *American Midland Naturalist* 73:225. | Fig. 3 | NO | We limited the re-analysis to 10- and 20-year intervals, 50-year intervals in this analysis. | NA | NA |

**Table S2. Segmented regression results.** NAs for initial breakpoint correspond to the origin in 1600.

| Breakpoint Year | Breakpoint Std. Error | Slope | Slope Std. Error |
|---|---|---|---|
| **1 Breakpoint (AIC -194.38)** | | | |
| NA | NA | 0.000384 | 0.00010231 |
| 1920 | 27.08705 | -0.0020588 | 0.0016684 |
| **2 Breakpoints (AIC -206.39)** | | | |
| NA | NA | -2.22E-04 | 0.0002009 |
| 1803.76 | 22.80658 | 1.62E-03 | 0.00056531 |
| 1911.453 | 10.92632 | -2.99E-03 | 0.0010104 |
| **3 Breakpoints (AIC -204.49)** | | | |
| NA | NA | 0.00098869 | 0.0016886 |
| 1660 | 46.82372 | -0.0006208 | 0.00037871 |
| 1790.548 | 18.53322 | 0.0016221 | 0.00042998 |
| 1908.776 | 11.22851 | -0.002791 | 0.0010103 |
| **4 Breakpoints (AIC -203.6067)** | | | |
| NA | NA | 0.0017809 | 0.0016866 |
| 1600 | 19.56001 | -0.0023037 | 0.0014848 |
| 1706.126 | 23.51232 | 0.00017891 | 0.00059719 |
| 1807.547 | 32.97915 | 0.0015919 | 0.00056462 |
| 1910.186 | 11.33187 | -0.0028803 | 0.0010092 |
